# Assessing heatdome effects on forest dynamics in the Pacific Northwest using Planet imagery

**DOI:** 10.64898/2026.01.12.699022

**Authors:** Aji John, Kavya Pradhan, Michael J. Case, Ailene K. Ettinger, Janneke Hille Ris Lambers

**Affiliations:** Department of Biology, University of Washington, Seattle, WA 98195, United States; The Nature Conservancy, 74 Wall Street, Seattle, WA 98121, United States; Plant Ecology, Institute of Integrative Biology, D-USYS, ETH Zürich, Switzerland

**Keywords:** heatdome, forest resilience, canopy density, PNW

## Abstract

Increasingly frequent extreme heat events are imposing significant stress on forest ecosystems, with implications for forests and the organisms that live in them. This study evaluates the effects of the June 2021 Pacific Northwest (PNW) heatdome (a prolonged high-pressure system that generated record-breaking temperatures) on forest canopy health and its subsequent recovery within the conifer-dominated Ellsworth Creek Preserve in Washington state, USA, using high-resolution Planet Labs satellite imagery. We analyzed normalized difference vegetation index (NDVI) trends from spring to fall from 2017 to 2024 to identify canopy stress and quantify changes in canopy greenness. Experimental forest restoration thinning treatments implemented to test their effectiveness in restoring old-growth forest characteristics at our sites allowed us to assess impacts of the heatdome event relative to these restoration treatments. Our results show a marked decline in summer NDVI following the 2021 heatdome, signaling substantial canopy stress. This decline, different than patterns in the other seven years, was particularly strong in dense, unmanaged (control) forest stands, which exhibited more pronounced NDVI declines than thinned (treatment) stands. These more pronounced stress signals during and after the heat event were associated with greater tree canopy height, canopy complexity, and tree density. The summer NDVI trends over the three post-heatdome years suggest a gradual recovery in tree canopy greenness, with thinned stands showing faster NDVI recovery than controls. In all, these results suggest that restoration thinning may improve canopy resilience to extreme heat. However, recovery trajectories varied widely within stand types, with persistent evidence of stress in high-density areas that suggest incomplete recovery as of 2023. Broadly, these findings show the vulnerability of dense forest canopies to extreme heat events, suggesting that forests with high canopy density may be less adaptable to recurring heat events. Nonetheless, managing forests through restoration thinning may enhance forest resilience to these climate-induced stressors – providing potential guidance for adaptive management strategies to maintain forest health in a world with more extreme climates.

## Introduction

Forests provide critical ecosystem functions, including the uptake of carbon, harboring of understory biodiversity, provisioning of wood products and cultural and recreational values (Trumbore et al. 2015). The long term ability of forests to provide these ecosystem functions depends on their ability to withstand the impacts of climate change (Kirilenko and Sedjo 2007, Seidl et al. 2017, Hartmann et al. 2022), as rising temperatures and changing precipitation regimes increases the vulnerability of most forests (Knapp et al. 2015). For example, hotter droughts have been linked to higher than background levels of tree mortality (Salomón et al. 2022), with these negative effects projected to worsen under future climate change in the Pacific Northwest (PNW) region (Mote and Salathé 2010, Diffenbaugh et al. 2015).

The long-term effects of rising temperatures on forests are thought to be negative, with short-term extreme heat events, like heat waves and heatdomes, likely to have additional negative impacts. These extreme events are often only 3-5 days, and long-lived trees generally survive them. However, heatwaves can increase stem dehydration in conifers (Salomón et al. 2022) and reduce radial growth (Pichler and Oberhuber 2007). Heatwaves can also exacerbate impacts of longer-term drought leading to even greater tree mortality (Van Mantgem et al. 2009, Allen et al. 2010, Perkins-Kirkpatrick and Lewis 2020). The timing of the extreme heat events, along with legacy effects of preceding seasons, may add complexity to the effects of heatdomes on forests (Salomón et al. 2022). Although there is still much we do not know about how forests are affected by extreme heat events, which have historically been relatively rare in the PNW, their increased incidence in the last few decades has resulted in increased interest in their long-term impacts on forest canopy health (Asner et al. 2016, Bastos et al. 2020, Anderegg et al. 2020).

While our understanding of impacts of short-term extreme heat events on forests is increasing, a significant limitation is that assessments of heatdome impacts must be opportunistic, as the timing of relatively rare heat waves and heatdomes are hard to predict. Continuously collected remote sensing data may thus provide a valuable opportunity to assess impacts of heat waves on forests after they have occurred. Remote sensing data have already been useful in assessing tree and forest mortality events (Foster et al. 2017, Hartmann et al. 2022) and demonstrate long-term impacts of climate change on forest health (Franklin et al. 2000, Senf et al. 2020), as well as to detect forest recovery from large disturbances (Brodrick and Asner 2017). Whether these data can be effective in measuring impacts and recovery from short-term heat events is not well known (but see (Sibley et al. 2025)). However, if heatwaves and heatdomes affect trees substantially enough to provide optical signals of heat stress via needle browning and desiccation, impacts of heatwaves on forest health may be observable and quantifiable using remote sensing based indices of tree health (like the commonly used Normalized Difference Vegetation Index (NDVI) (Breshears et al. 2005). If so, these approaches can be used to measure both short-term impacts and long-term recovery of forests following such events.

To explore the impacts of extreme events on the health of forest ecosystems, measured by remote-sensing techniques, we assess how a conifer-dominated coastal forest in the Pacific Northwest of USA was impacted by an extreme heat event in 2021 (White et al. 2023). The extreme heat event was a ‘heatdome’, a meteorological phenomenon in which a high-pressure system trap warm air over a region, leading to prolonged periods of extreme heat. We take advantage of an existing long-term forest restoration experiment, which applies different forest density treatments, to explore how canopy density influences short-term impacts and long-term recovery (Case et al. 2021, 2023, Chamberlain et al. 2021). We ask three key questions about how these forests responded to the 2021 Pacific Northwest (PNW) heatdome. First, we ask whether canopy damage can be detected using high-resolution remotely sensed imagery, comparing NDVI trends in the heatdome year with those in seven other more ‘normal’ years. We expect that heat stress during the heatdome would result in visible vegetation changes, realized as a decrease in NDVI associated with leaf desiccation. Second, we ask what vegetation characteristics contribute to increased canopy stress during and immediately after the heatdome (Fig. 1), taking advantage of the restoration treatments imposed in these forests. We expect that denser canopies would experience more severe stress, due to their greater thermal mass and potential to trap heat. Finally, we ask whether forest canopies show evidence of recovery in the years following the heatdome event, and whether restoration treatments affect the pace of recovery. We expect that NDVI values would gradually increase over time after the event, indicating recovery, but that the pace and extent of recovery would vary based on initial canopy structure and treatment history. Additionally, we compared alternative model structures to evaluate which best captured seasonal NDVI dynamics and canopy recovery following the 2021 heatdome.

**Figure 1.**
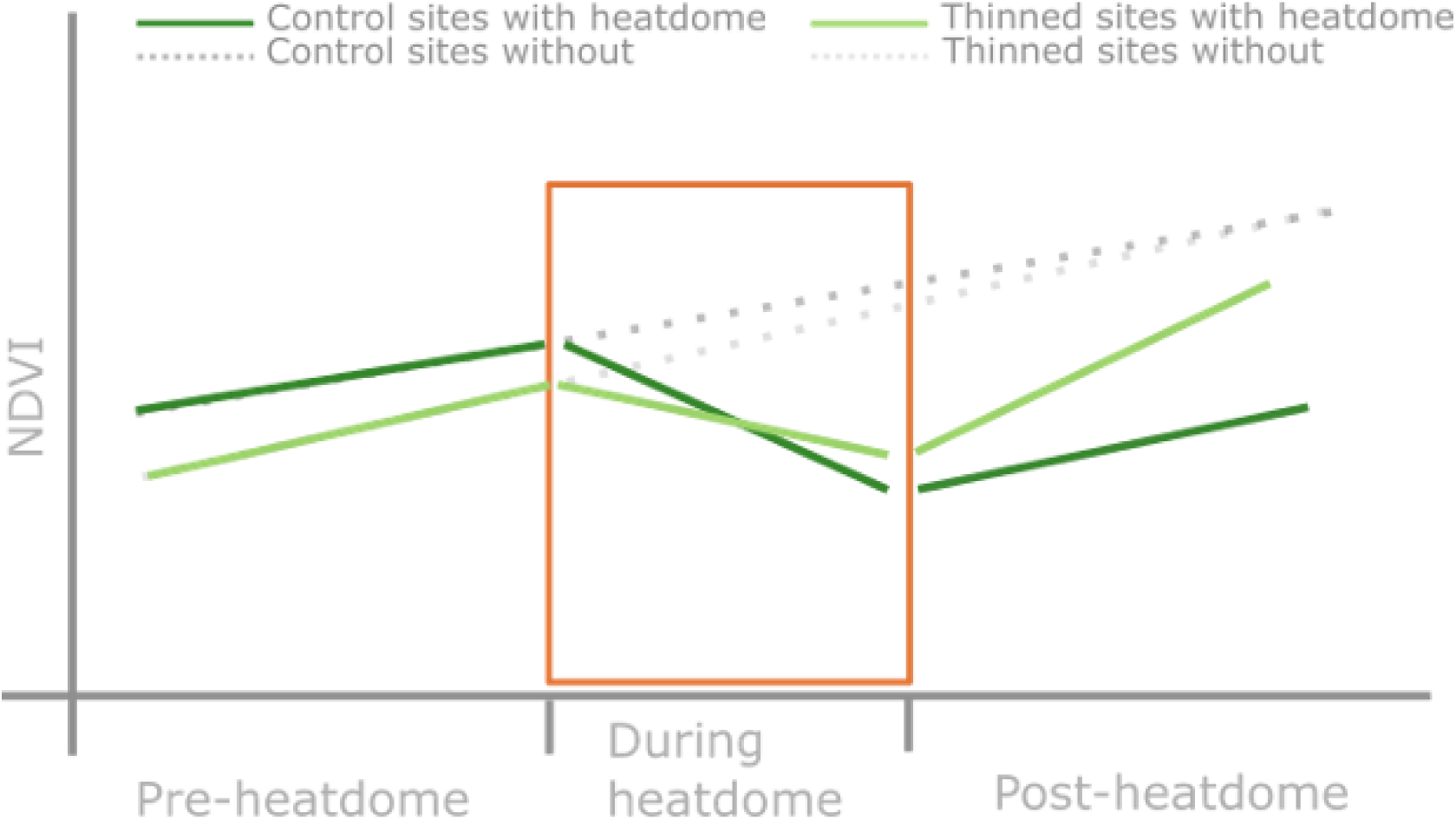
Hypothesized relationships between forest canopy cover and remotely sensed normalized difference vegetation index (NDVI). In the absence of a heatdome event, NDVI (representing greenness of forest canopy) values in coniferous forests are expected to be stable (grey). When an extreme heat event occurs, we expect NDVI to decrease with greater decreases in more dense forests (dark green) compared to thinned forests (light green). Over time, the NDVI in these forests will recover as time from heatdome increases as demonstrated by the increasing NDVI values since heatdome.

## Materials and Methods

### Study area

We conducted this study in a preserve located in the Pacific Northwest Coastal Ecoregion (Dinerstein et al. 2017), characterized by conifer dominated forests, large swaths of which have been heavily logged and converted to timberlands. Our sites were located at Ellsworth Creek Preserve (Ellsworth) in the Willapa Hills (WA, USA), a 33 km^2^ forested preserve, in which large parts were active timberlands under private industrial ownership until the late 1990s (Fig. 2). Ellsworth is now managed by The Nature Conservancy (TNC) and is part of a watershed-level experiment aimed at restoring old-growth forest characteristics to former timberlands. Like other forests in this ecoregion, coastal mixed-conifer rainforests dominate and experience warm-dry summers and cool-wet winters. The preserve, which includes a mix of second growth and remnant old-growth stands, was established as a watershed scale experiment, with sub-basins in the Ellsworth Creek watershed allocated to two main forest management types: thinning forest stands or leaving stands without intervention (as control). Further details on the ongoing experiment can be found in Case et al. (2023).

**Figure 2:**
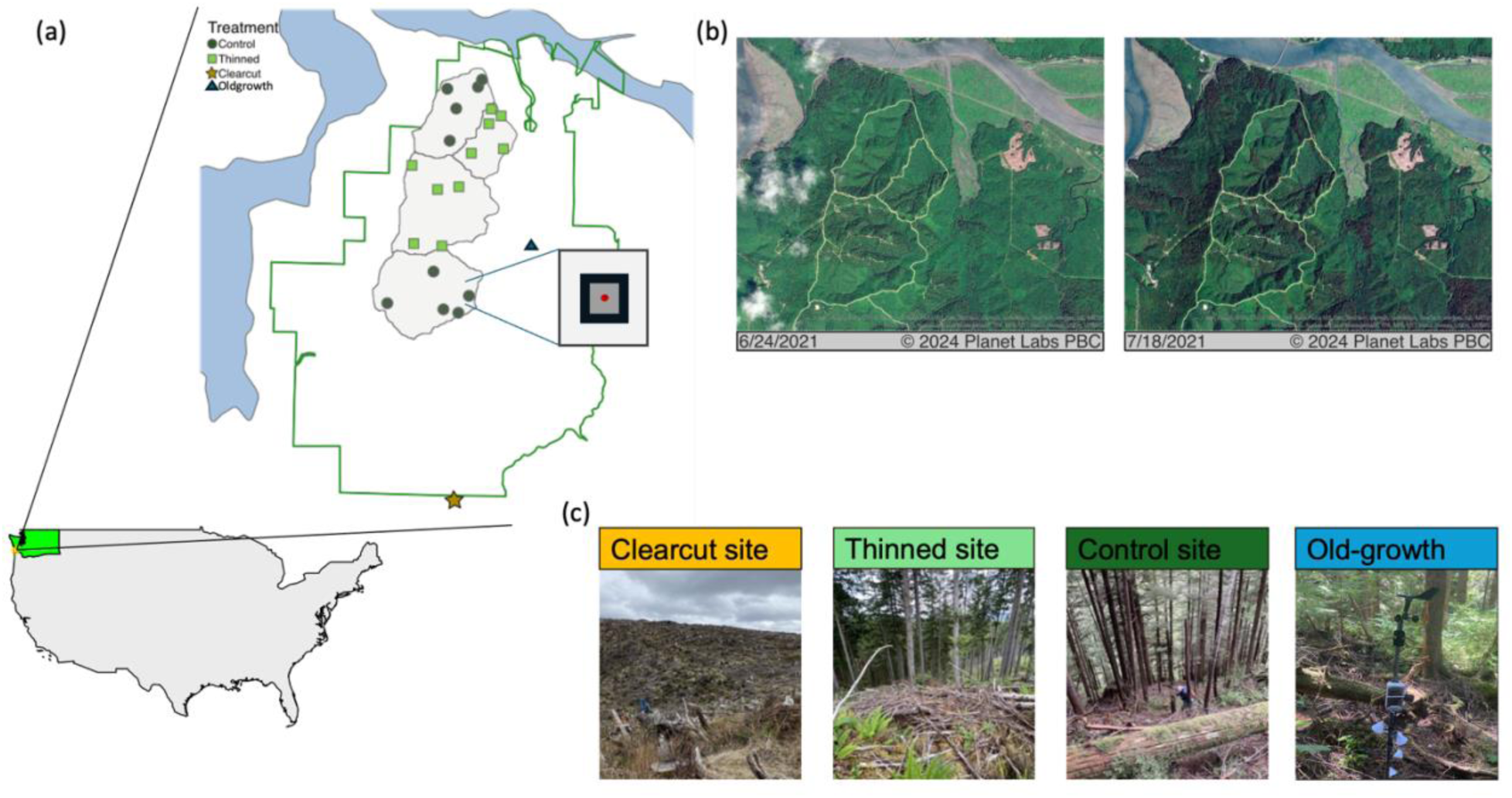
Study site and experimental design. (a) Location of the study region (Ellsworth Creek Preserve in the Willapa Hills, Washington State) in the PNW of United States (inset) and distribution of treatment sites within the study landscape, including clearcut, thinned, control, and old-growth stands. For each site, we extracted NDVI data from Planet by averaging the 3 × 3 cells surrounding the focal pixel in which our sites were located to obtain final NDVI values. (b) Representative Planet imagery illustrating forest conditions before and after the 2021 heatdome event. (c) Ground-level photographs showing typical conditions at each treatment type.

### Remote sensing data

We examined the short-term and long-term impacts of the 2021 extreme heat event on forest canopy using remote sensing-based metrics from Planet Labs. Planet Labs (Planet 2018) operates a fleet of satellites called Super Doves which image the earth daily, giving a ground sample distance (physical distance on the ground represented by each pixel) of 3-5m. PlanetScope, a 4-band level-2 product that gives surface reflectance was downloaded and prepared for analysis using a cloud-agnostic workflow execution provider called SWEEP (John et al. 2019). Prior to applying restoration treatments, TNC established permanent vegetation monitoring plots (17.8m radius circular plots;) within all sub-basins. We monitored a subset of these plots (20 plots; ten in un-thinned (control) and ten in thinned stands) in the North and Central sub-basins selected based on topographic position index (refer to (John et al. 2024) for details). One plot in a clear-cut and one in an old-growth remnant were included for additional comparison.

For the 22 plots, we downloaded imagery for 8 years (2017-2024) and clipped to a 30m x 30m-pixel area (∼ 900 m square area from the center of the plot). From each clipped image, we extracted spectral summaries including the minimum, maximum, and mean values for the red, green, blue (RGB), and near-infrared (NIR) bands, as well as the NDVI. Imagery with more than 15% cloud cover was not included in the analysis. We used categorical variables as a proxy for canopy density (Clear Cut = no canopy, Old-Growth = very dense canopy, Thinned = less dense canopy and Control = moderately dense canopy) as their relationship to canopy cover has been established by previous studies conducted in this experimental forest (Chamberlain et al. 2021, Case et al. 2023).

### Statistical Analysis

All statistical analyses were conducted in R version 4.1.2 (R Core Team 2021). Data wrangling and preprocessing were carried out using the *tidyverse* package (Wickham et al. 2019). We evaluated (1) the detectability of canopy stress during the 2021 Pacific Northwest heatdome using remotely sensed NDVI, (2) the influence of forest structure and treatment history on heat-related NDVI suppression, and (3) recovery trajectories in subsequent years. Because NDVI trends are subtle and non-linear, we used multiple modeling approaches to explore our data, including linear regressions, linear mixed-effect models (LMMs) and GAMs. This allowed us to capture both linear and nonlinear seasonal trends, while accounting for repeated measurements at the same sites and improving robustness by comparing results across models (Gelman and Hill 2006).

To address our first question, whether there is anomaly in 2021 trends compared to the baseline seasonal trends, we first fit year-specific linear models estimating the relationship between NDVI and day of year (DOY) from 2017 through 2024. DOY was defined as the calendar day beginning on January 1 (DOY = 1) and ending on December 31 (DOY = 365 or 366 in leap years). These models provided annual estimates of NDVI seasonal directionality, defined as the overall trajectory of NDVI within each year — whether NDVI increased (greening), decreased (suppression), or remained stable over the season. Directionality was quantified using correlation coefficients and linear slopes. We classified trends as suppression (R < –0.2), greening (R > 0.2), or stable (–0.2 ≤ R ≤ 0.2), based on thresholds for moderate correlation strength in ecological time series (White et al. 2009, Pompa-García et al. 2021a, Guo et al. 2025). To assess structural recovery and canopy persistence following the 2021 heatdome, we analyzed the minimum NDVI (NDVIₘᵢₙ) within each growing season. NDVIₘᵢₙ captures the lower envelope of canopy greenness and reflects baseline foliage retention. Also, for seasonal trajectory analyses, we evaluated maximum NDVI (NDVIₘₐₓ) as an indicator of peak canopy greenness and stress intensity.

To address the second question on whether vegetation characteristics contributed to increased canopy stress, we fit a generalized additive model (GAM) using the *gam*() function in the *mgcv* package (Wood 2001) to capture non-linear seasonal NDVI dynamics. The response variable was NDVI, modeled as a smooth function of DOY with an interaction of forest treatment (a categorical variable – old-growth, clearcut, thinned, and control) and three temporal periods (Pre-2021, 2021, Post-2021). This model provided a more flexible estimation of seasonal patterns across varying forest types and heatdome intervals than linear models do. Model performance was evaluated using adjusted R², RMSE, AIC, and deviance explained. Residual diagnostics (*gam.check*) were used to verify the absence of heteroscedasticity, temporal autocorrelation, and over smoothing. Model fit was optimized using restricted maximum likelihood (REML). To quantify treatment and period-specific responses, we extracted marginal effects of DOY on NDVI from the fitted GAM smooths. These effects represent the local slopes of the seasonal NDVI trajectory, with positive values indicating greening (increasing NDVI with DOY) and negative values indicating suppression (declining NDVI with DOY). For each treatment × period group, we summarized the minimum and maximum marginal effects along with approximate 95% confidence intervals, which allowed us to identify whether seasonal NDVI changes were consistently positive, negative, or indistinguishable from zero. Furthermore, to evaluate how the rate of canopy greenness changed throughout the growing season, we derived partial effects of DOY from the fitted GAM model. Partial effects give us the contribution of DOY to NDVI variation after accounting for forest treatment and temporal period effects, thus providing shape of the seasonal NDVI trajectory.

Finally, to address the third question on evidence of recovery and the role of canopy density on rates of recovery, we tested statistical differences in NDVI values while accounting for repeated measures within sites, by fiting a linear mixed-effects model (LMM) using the *lmer*() function from the lme4 package (Bates et al. 2014). NDVI was the response variable, with DOY, treatment, period, and their interactions specified as fixed effects. Plot identity was included as a random intercept to control for non-independence of observations within plots. We used the *emmeans* package (Lenth 2025) to get estimated marginal means and to perform pairwise contrasts between treatment-period combinations. This enabled formal tests of differences in NDVI among groups such as old-growth forests before and after the heatdome (i.e. OLD_GROWTH:Post2021 vs Pre2021).

To address potential concerns of using varying modeling approaches, we compared the relative fit of linear, GAM, and mixed-effects models using a common set of metrics (adjusted R², RMSE, AIC, deviance explained) and is reported in the Results section.

## Results

Overall, we found evidence that the 2021 Pacific Northwest heatdome caused significant reductions in forest canopy greenness, with impacts varying across restoration treatments. NDVI seasonal trajectories showed reduced greening in 2021 following the heatdome, particularly in old-growth and denser canopy stands. Sites with lower canopy density (thinned sites) appeared to be less impacted by the heatdome and exhibited faster recovery than control sites. Here, we describe our results relative to the specific questions we had.

### Does forest canopy show signs of stress following an extreme heat event?

The seasonal NDVI trajectories revealed clear differences across years and treatments, reflecting both the immediate impacts of the 2021 heatdome and subsequent patterns of canopy stress and recovery (Fig. 3, Table 1). In 2021, we observed a negative correlation between NDVI and day of year (DOY) (R = −0.22, p < 0.05), indicating a decline in canopy greenness over the growing season. All other years either showed stable seasonal trajectories or positive greening trends, particularly 2022 (R = 0.34) and 2024 (R = 0.21). NDVI trajectories in 2021 diverged markedly from previous years.

**Figure 3.**
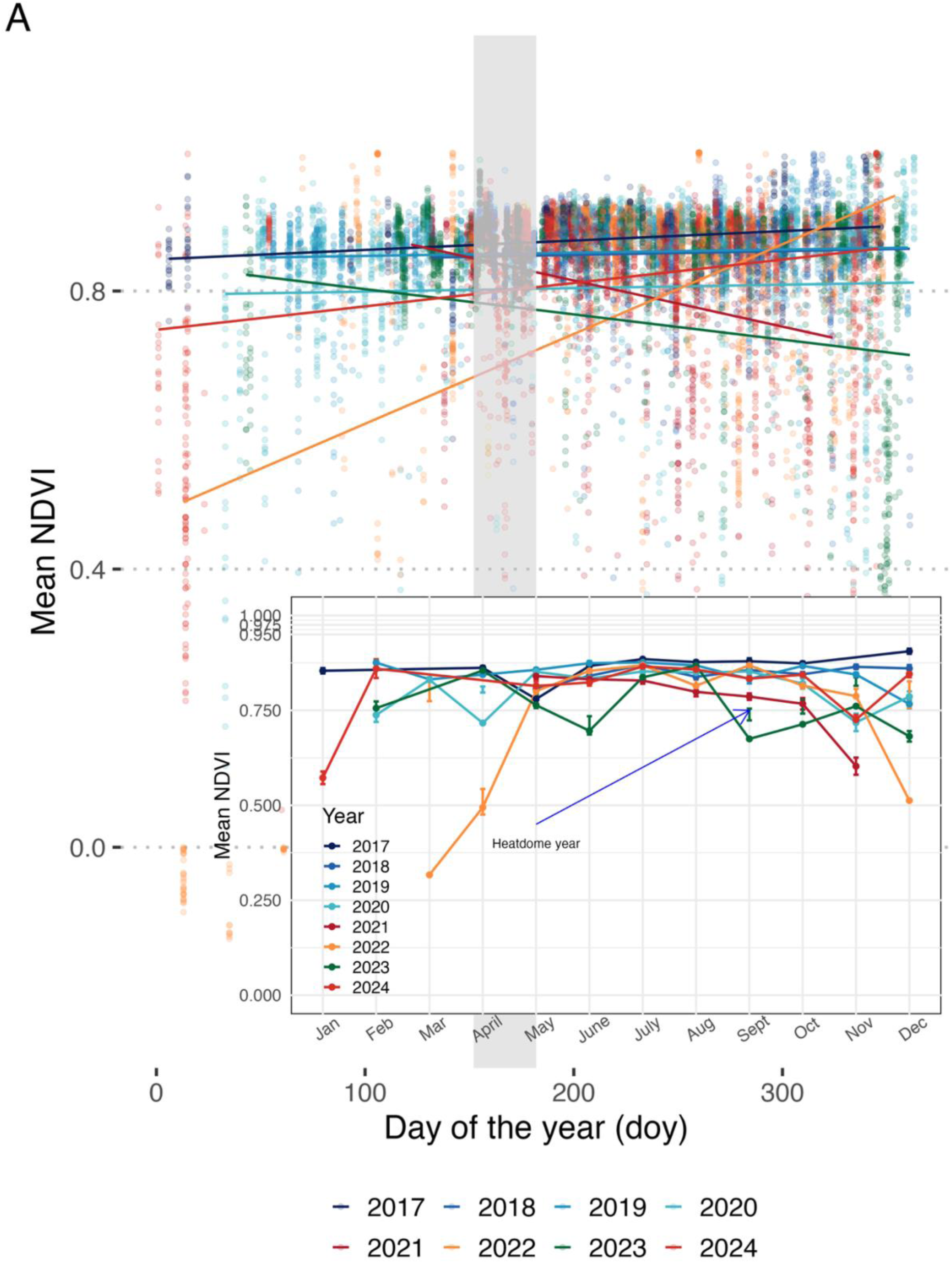
Seasonal NDVI in forests before, during and after the 2021 PNW heatdome event. The dots refer to the plot level NDVI (closer to 1 indicates healthier vegetation). Mean NDVI values regressed against day of the year and inset shows the values by month (A). The smaller number of points in 2017 is because of limited availability of imagery in the early years of Planet product release, and absence of points in first few months of 2018 and in first and last few months of 2021 is because of scene quality and cloud filter. For all years, the rectangle refers to June and is shaded in the heatdome year (2021).

**Table 1:** Year-wise correlation between NDVI and day of year (DOY), including slope estimates, p-values, and classified trend interpretations. Analysis used data filtered for cloud cover less than 15%.

| Year | Correlation coefficient (R) | Slope | p-value | Trend |
| --- | --- | --- | --- | --- |
| 2024 | 0.213 | 0.000358 | <0.001 | Greening |
| 2023 | -0.096 | -0.000330 | <0.001 | Stable |
| 2022 | 0.340 | 0.001268 | <0.001 | Greening |
| 2021* | -0.222 | -0.000670 | <0.001 | Suppression |
| 2020 | 0.020 | 0.000038 | 0.414 | Stable |
| 2019 | 0.038 | 0.000052 | 0.128 | Stable |
| 2018 | 0.056 | 0.000130 | 0.036 | Stable |
| 2017 | 0.086 | 0.000129 | 0.016 | Stable |

The GAM model explained a moderate proportion of variance (adjusted R² = 0.19, RMSE = 0.176, AIC = –8731), and residual diagnostics (*gam.check*) showed no evidence of temporal autocorrelation or heteroscedasticity. Model predictions indicated suppressed NDVI in old-growth and control plots in 2021 (the heatdome year), where seasonal NDVI peaks were flattened (Fig. 4). This pattern contrasts with expected seasonal greening observed in pre-2021 years. The GAM also revealed significant treatment × period effects (p < 0.001 across multiple smoothing terms; Table 2), confirming that seasonal NDVI trajectories differed between forest types and across time periods. Smooth terms for all treatment × period combinations were statistically significant (p < 0.005), with effective degrees of freedom (EDF) ranging from 1.01 to 8.49, indicating moderately non-linear seasonal patterns that differed among treatments. However, differences between forest types during and after the heatdome were noticeable. For example, old-growth stands in 2021 had an EDF of 1.01 and a moderate F-value of 2.98 (p = 0.081). Control plots in 2021 showed a pronounced seasonal NDVI pattern, with an EDF of 1.77 and an F-value of 15.30 (p < 0.001). Denser canopy plots, particularly control stands, exhibited evidence of seasonal NDVI suppression following the heat dome, with seasonal NDVI patterns evident in both the 2021 and Post-2021 periods. In contrast, thinned plots showed comparatively less pronounced changes in seasonal NDVI patterns during 2021. Although thinned plots also exhibited detectable seasonal NDVI variation (p ≤ 0.005), the magnitude of seasonal change was lower than that observed in control stands. In 2021, the heat dome year, thinned plots had an EDF of 1.89 and an F-value of 6.15, which was lower than the corresponding control values (EDF = 1.77, F = 15.30) and comparable to old-growth (EDF = 1.01, F = 2.98). Similarly, during the Post-2021 period, the F-value for thinned plots was 72.59, while old-growth and clearcut plots had F-values of 13.17 and 10.84, respectively, reflecting variation in seasonal NDVI among stand types.

**Figure 4.**
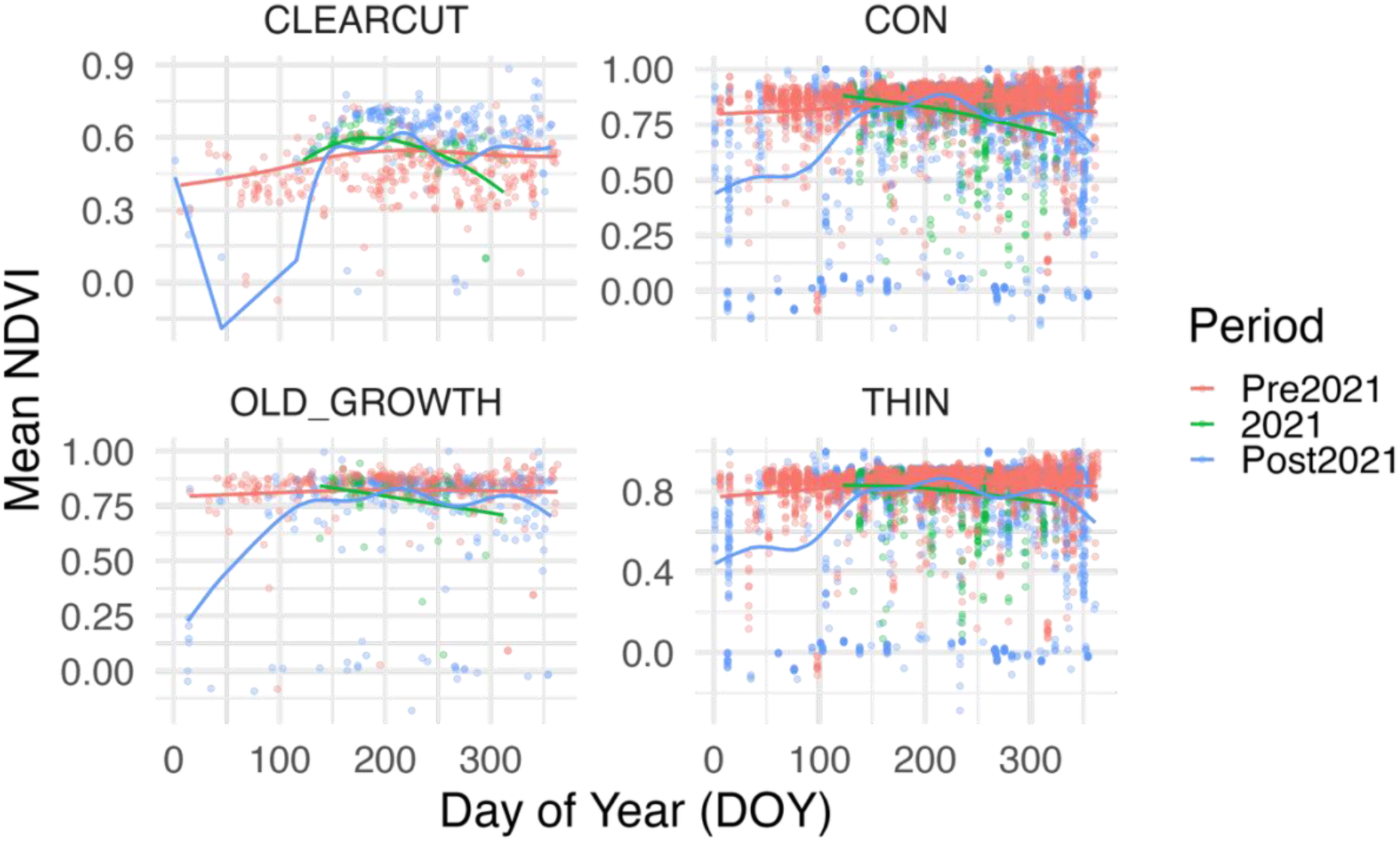
Seasonal NDVI trajectories by treatment and period based on GAM predictions. Points show observed NDVI, and smooth colored lines represent fitted seasonal trends for each temporal period (Pre-2021, 2021 heatdome, and Post-2021). Each panel shows a different forest treatment (clearcut, control, old-growth, thinned). During 2021, old-growth and control plots show flatter seasonal curves indicating heat-related canopy stress, while thinned plots exhibit relatively stable NDVI dynamics, suggesting improved resilience and partial recovery after the heatdome.

**Table 2:**
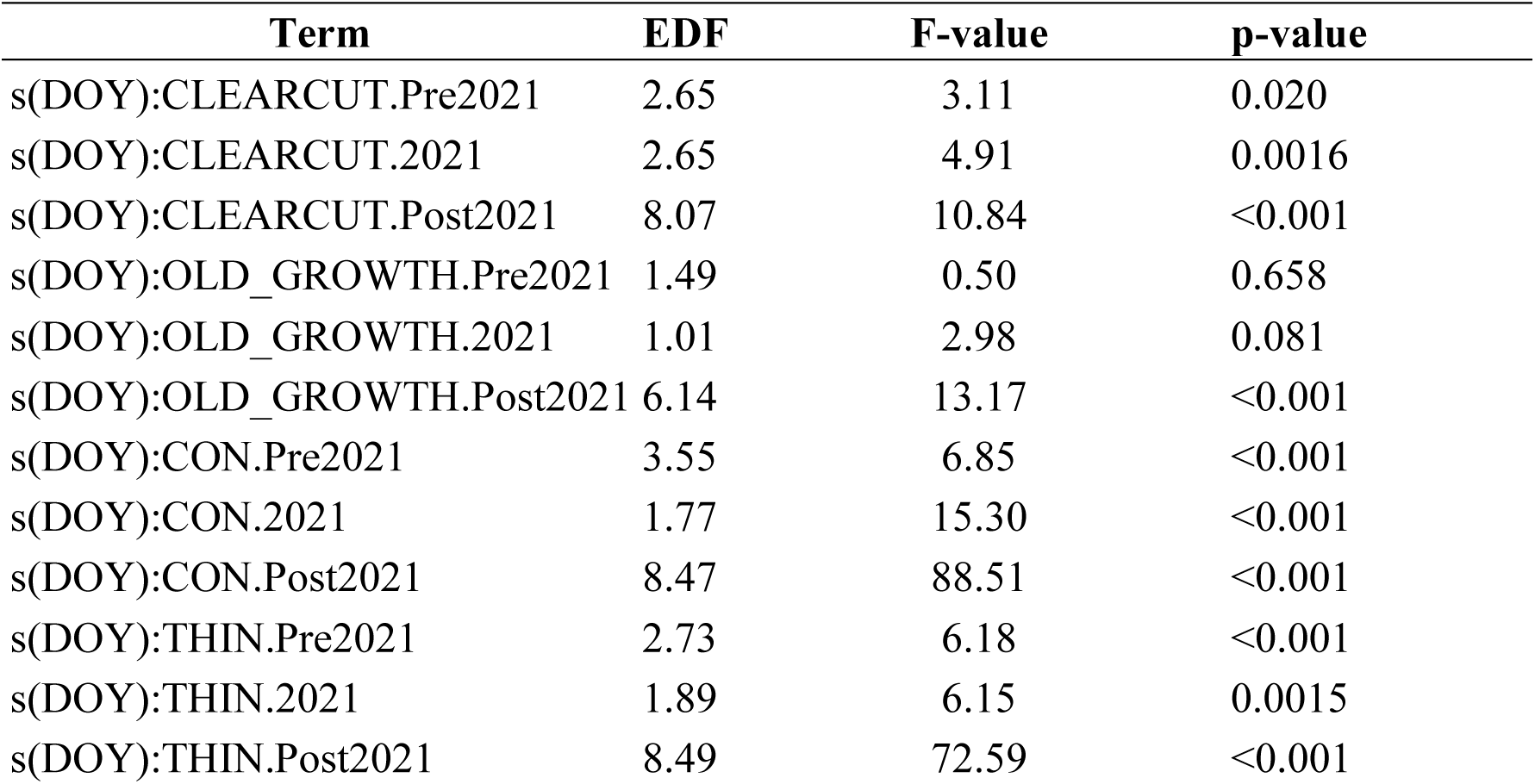
Summary of GAM smooth terms for NDVI ∼ DOY where each row is the smooth term for DOY within a treatment × period group. Values shown are effective degrees of freedom (EDF), F-statistics, and p-values for each smooth term. EDF relates to model complexity of the smoothing term; ∼1 signifies linear pattern, and greater than 2 means non-linear pattern. The F-value and p-value signify whether the seasonal curve is statistically different from flat (no seasonal pattern).

| Term | EDF | F-value | p-value |
| --- | --- | --- | --- |
| s(DOY):CLEARCUT.Pre2021 | 2.65 | 3.11 | 0.020 |
| s(DOY):CLEARCUT.2021 | 2.65 | 4.91 | 0.0016 |
| s(DOY):CLEARCUT.Post2021 | 8.07 | 10.84 | <0.001 |
| s(DOY):OLD_GROWTH.Pre2021 | 1.49 | 0.50 | 0.658 |
| s(DOY):OLD_GROWTH.2021 | 1.01 | 2.98 | 0.081 |
| s(DOY):OLD_GROWTH.Post2021 | 6.14 | 13.17 | <0.001 |
| s(DOY):CON.Pre2021 | 3.55 | 6.85 | <0.001 |
| s(DOY):CON.2021 | 1.77 | 15.30 | <0.001 |
| s(DOY):CON.Post2021 | 8.47 | 88.51 | <0.001 |
| s(DOY):THIN.Pre2021 | 2.73 | 6.18 | <0.001 |
| s(DOY):THIN.2021 | 1.89 | 6.15 | 0.0015 |
| s(DOY):THIN.Post2021 | 8.49 | 72.59 | <0.001 |

The first derivatives of the GAM smooths, representing the marginal effects of day of year (DOY) on NDVI, showed distinct seasonal variation across treatments and periods (Table A2; Fig. A1). During the 2021 heatdome, mean slopes were consistently negative in old-growth, control, and thinned plots (–0.00035 to –0.00088), indicating a mid-season suppression of greenness relative to pre-2021 patterns. The largest decline occurred in control plots (–0.00088). By contrast, in Post-2021, the mean slopes for all treatments became positive, particularly in old-growth (0.00148) and thinned (0.00054). Clearcut and thinned treatments also exhibited the widest range of slopes (–0.015 to +0.022). The Pre-2021 period shows small but consistently positive slopes across treatments (on the order of 10⁻⁴). Overall, these marginal effects confirm that 2021 was characterized by a pronounced reduction in NDVI rate of change (i.e., greening slowdown), followed by compensatory recovery by 2023–2024.

NDVIₘₐₓ declined across all treatments during the 2021 heatdome year (Fig. A2b). The decline was most pronounced in old-growth and control plots. By 2023–2024, thinned and control stands showed partial rebounds in NDVIₘₐₓ (approximately 0.86–0.88 in 2021 to ∼0.82–0.90 in the Post-2021 period), whereas old-growth and clearcut plots remained below pre-2021 levels. These patterns confirm the GAM-derived signal of heat-induced canopy suppression.

### What is the trajectory of forest recovery from an extreme heat event?

Overall, most forests began to recover after the 2021 heat event, but the speed and consistency of recovery depended on how open the canopy was. Partial effects of DOY on NDVI estimated from the GAM (Fig. 5) characterized the seasonal greenness responses that varied across treatments and time periods. Before 2021, effects were near zero for all treatments, indicating stable intra-annual NDVI dynamics. During the heatdome period, partial effects became negative mid-season, with the strongest declines in old-growth and control stands (minimum ∼ –0.10 to – 0.14 NDVI day⁻¹), reflecting suppressed greening. In contrast, thinned and clearcut stands showed weaker and short-lived declines, with slopes remaining close to zero. By 2023 (Post-2021), partial effects returned toward zero or slightly positive values (mean ∼ 0.001–0.005), although variability increased, particularly in open-canopy stands (clearcut > thinned).

**Figure 5.**
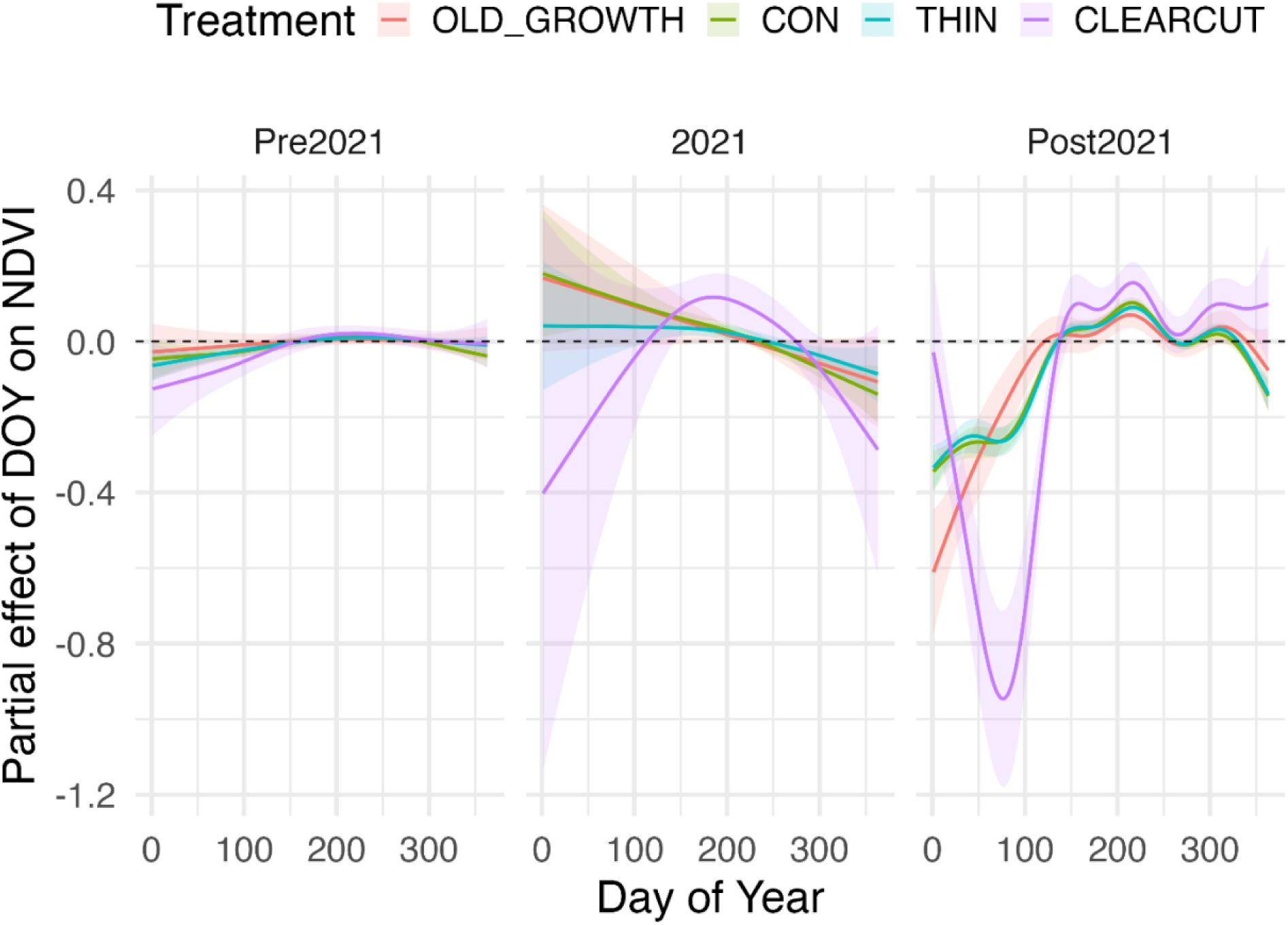
Partial effects of DOY on NDVI across forest treatments and periods from the GAM. Partial effect curves show the contribution of DOY to NDVI variation after accounting for treatment and period effects. Lines represent fitted partial effects and shaded ribbons indicate 95% confidence intervals across three temporal periods: Pre-2021, 2021 (heatdome year), and Post-2021. During 2021, particularly in CON and OLD_GROWTH plots, partial effects were negative in mid-season, indicating suppression of greening during the heatdome. By contrast, thinned and clearcut plots maintained positive partial effects across periods.

Recovery trajectories of old-growth showed evidence of partial, but incomplete NDVI rebound, smooths from 2022 and 2023 remained below pre-2021 baselines, and marginal effects of DOY were weaker (Fig. 4). In control Post-2021, effects remained near zero. Also, thinned and clearcut plots showed no evidence of delayed recovery. While overall NDVI values were lower in clearcut, these plots did not exhibit further suppression post-2021, and marginal effects gradually returned to positive values by 2024. The strongest NDVI suppression was observed in clearcut and control plots during and immediately following the 2021 event. Marginal effects of DOY on NDVI were highly negative in these treatments (Table A2, Fig. 5), ranging from –1.27 to –3.93, with 95% confidence intervals consistently below zero across large portions of the growing season. In contrast, thinned plots exhibited relatively stable NDVI responses across all periods, with marginal DOY effects remaining weakly positive and confidence intervals overlapping zero. Old-growth plots displayed intermediate patterns, with some suppression during 2021 but evidence of partial seasonal recovery in Post-2021 periods.

The LMM supported the GAM-derived recovery trajectories, providing complementary evidence that forest canopies showed both immediate heat stress and gradual recovery following the 2021 heatdome. Fixed effects revealed significant NDVI declines during 2021, followed by modest rebounds in 2022–2023 (Table 3; full LMM results can be found in Appendix Table A2). The model explained a moderate proportion of NDVI variation (conditional R² = 0.14; RMSE = 0.18). NDVI remained lower in the Post-2021 period relative to Pre-2021 (β = –0.24 to –0.30, *t* ≈ –4.3, *p* < 0.01). The negative DOY × Period2021 interaction (*t* = –2.36) captured the seasonal suppression of greenness during the heatdome, while continued negative interaction effects for THIN and CON treatments during Post-2021 (β ∼ –0.21 to –0.24) suggested incomplete recovery (Figure 6).

**Figure 6.**
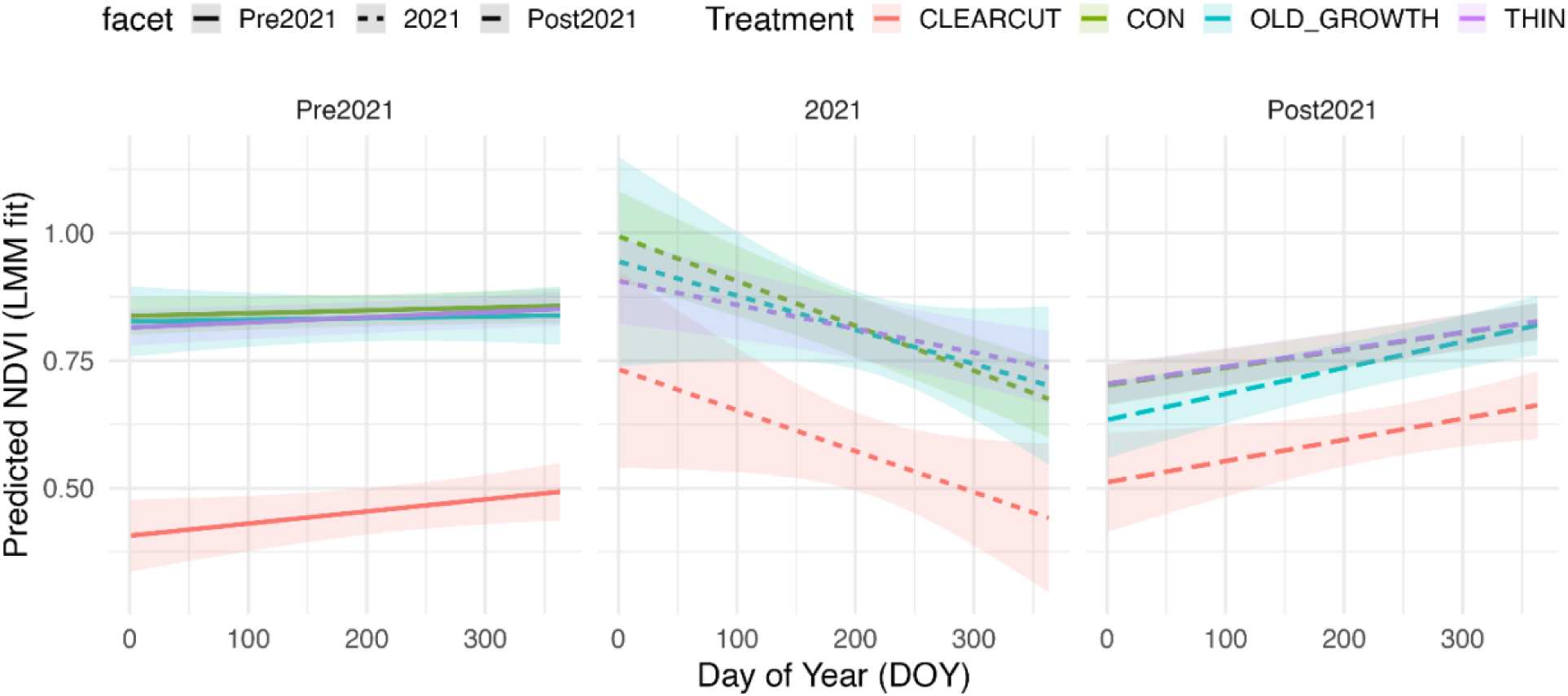
Predicted NDVI trajectories from the LMM across forest treatments and periods. Lines represent model-predicted NDVI as a function of day of year (DOY). Shaded ribbons indicate 95% confidence intervals. Across treatments, NDVI declined sharply during the 2021 heatdome, followed by gradual post-2021 increases. Thinned and control plots exhibited relatively stable or rebounding NDVI trends post-2021, while old-growth and clearcut sites showed slower or more variable recovery trajectories.

**Table 3:** Summary of LMM fixed effects for NDVI as a function of day of year (DOY), forest treatment, and period. Full detail of the LMM fixed effects model can be found in Appendix A Table A2.

| Effect | Estimate ($\beta$) | SE | *t* value |
| --- | --- | --- | --- |
| Intercept | 0.41 | 0.04 | 11.25 |
| DOY | 0.00024 | 0.00012 | 1.91 |
| Period (2021) | 0.33 | 0.10 | 3.16 |

| Effect | Estimate ($\beta$) | SE | $t$ value |
| --- | --- | --- | --- |
| Period (Post-2021) | 0.10 | 0.06 | 1.81 |
| DOY × Period (2021) | -0.0010 | 0.0004 | -2.36 |
| TRT (THIN × Post-2021) | -0.22 | 0.06 | -3.89 |
| TRT (CON × Post-2021) | -0.24 | 0.06 | -4.35 |

Values of NDVIₘᵢₙ remained depressed in old-growth and control plots through 2022 (Fig. A2a). In contrast, thinned and clearcut plots exhibited gradual increases in NDVIₘᵢₙ across subsequent years. These results complement the GAM-modeled seasonal trajectories, showing that while greenness peaks remained subdued, baseline canopy greenness gradually increased post-heatdome.

## Model fits across approaches

Across all approaches, model fit improved when hierarchical structure was included (Table 4). Year-specific linear models explained relatively little variance (mean R² = 0.03, RMSE = 0.167), whereas the LMM model improved performance (adj. R² = 0.11, RMSE = 0.183). The GAM provided the best overall representation of seasonal NDVI trajectories (adj. R² = 0.19, RMSE = 0.176, deviance explained = 19.3%), although with slightly higher RMSE due to its flexibility in capturing non-linear patterns. Together, our findings indicate that the GAM is most effective for capturing seasonal canopy trajectories dynamics across treatments and time periods.

**Table 4:** Comparison of model performance metrics across linear mixed-effects models (LMM), year-specific linear models, and generalized additive models (GAM) for NDVI ∼ DOY. Metrics shown are adjusted R², RMSE, AIC, and deviance explained.

| Model type | Adjusted $R^2$ | RMSE | AIC | Deviance explained |
| --- | --- | --- | --- | --- |
| Mean across years | 0.03 | 0.167 | -1224.4 | 2.9% |
| Linear mixed-effects model | 0.11 | 0.183 | -7472 | — |
| Generalized additive model | **0.19** | 0.176 | -8731 | 19.6% |

## Discussion

### Forest canopy is under stress following extreme heat

Our results demonstrate that forest canopy greenness, as measured by NDVI, was substantially impacted by the 2021 Pacific Northwest heatdome, declining by ∼10% (Fig. 3, Fig. 4 and Fig. 5). NDVI trajectories within 2021 showed a marked mid-season decline (following the early season heatdome) relative to previous years, indicating acute canopy stress and reduced overstory tree health under extreme thermal conditions. These patterns are consistent with canopy-scale thermal desiccation resulting from sustained heat exposure. In contrast, preceding years such as 2018 exhibited positive correlations between day of year (DOY) and NDVI (Fig. 3), reflecting regular seasonal greening in the absence of severe climatic disturbances. Seasonal NDVI patterns remained relatively stable across 2018 to 2020, suggesting that under normal conditions, forest canopy dynamics were resilient to typical interannual variability. Of course, year to year variability likely also influenced these trends, with the slight negative trend in NDVI in 2023 (Fig. 3). This likely reflects both the continuation of unusually low canopy greenness observed in 2022 following the 2021 heatdome, and additional heat stress associated with warmer than usual temperatures during the 2023 growing season (NOAA National Centers for Environmental Information 2023).

The NDVI response we observed following the 2021 heatdome are consistent with other studies examining effects of extreme heat events on forest canopy (Pettorelli et al. 2005, Yuhas and Scuderi 2009, Sang and Hamann 2023, Sibley et al. 2025). However, we also found that the magnitude of canopy stress varied across treatment types, with control and clearcut plots showing a greater decrease in NDVI compared to thinned plots during the 2021 heatdome (Fig. 4 and 5). These results suggest that forests with very dense canopy cover (control plots) are more vulnerable to heat stress during extreme heat events, whereas moderately thinned stands exhibited greater resilience. A caveat is that NDVI values may be influenced by environmental forcing like precipitation (Santos and Negri 1997). Coastal coniferous forests like the ones in our study are in a high precipitation zone and have a long growing season; declining NDVI in denser canopies following the heatdome might have been complicated by saturation in optical signal caused by shading induced by denser canopies (Yoder and Waring 1994, Leblanc et al. 1997, Brodrick and Asner 2017). NDVI has also been shown to decline during multi-year droughts, highlighting that remotely sensed greenness indices reliably capture vegetation stress under both acute and chronic climatic extremes (Yoshida et al. 2015, Song et al. 2019, Pompa-García et al. 2021b). While we do not have evidence of drought in our study system during the 2017–2024 observation window, the congruence between our results and drought-focused studies suggests that NDVI responses to canopy stress can be generalized across different types of climatic disturbance.

Other studies of conifer responses to the 2021 PNW heatdome have shown crown discoloration, desiccation and loss of greenness (Klein et al. 2022, Still et al. 2023, Sang and Hamann 2023, Sibley et al. 2025). Moreover, in drought-impacted or semi-arid regions, increases in canopy temperatures have resulted in adult tree mortality (Scherrer et al. 2011, Guha et al. 2018). Whether canopy stress following the heatdome is indicative of longer-term increased tree mortality is unclear as such events have been extremely rare in the PNW. Nevertheless, extreme heat events are expected to increase in frequency in the Pacific Northwest (Chen et al. 2023), so the potential impacts of increased canopy stress on tree mortality following individual or multiple extreme heat events needs to be monitored. These episodic events will likely exacerbate the effects of long-term warming (and *vice versa*) through press and pulse dynamics (Harris et al. 2018) and have potential to cause damage beyond the effects of a single extreme event alone. Moreover, less dramatic effects on forests, including crown die-back following heatdome events, can also alter the forest’s buffering capacity (Ruthrof et al. 2016) and perhaps predispose individual trees to other stressors, such as insect and disease outbreaks. More broadly, the increased plant stress (as indicated by the decrease in NDVI, Fig. 3) that we observed after the 2021 heatdome has the potential to decrease the ability of forests to buffer the understory from extreme heat. This effect could be stressful for understory organisms already coping with increasing average temperatures and altered precipitation patterns due to climate change.

### Thinning in secondary forests reduces canopy stress and promotes recovery

We found that control plots that had not been thinned experienced more pronounced NDVI declines during and after the 2021 heatdome (Fig. 4). Thinned restoration plots, by contrast, exhibited a slightly less severe NDVI decline during the heat event and maintained more stable seasonal NDVI dynamics across the ensuing years. Our results support other studies that suggest thinning may mitigate heat stress impacts by reducing canopy density, potentially enhancing air circulation and lowering heat load within the canopy structure (Case et al. 2023). These thinned plots not only seemed to show greater resilience, but they also showed a faster trajectory toward recovery following the extreme heat event. Specifically, post-heatdome responses differed notably between control and thinned plots. We found thinned plots consistently maintained higher NDVI values and more stable seasonal patterns compared to control plots across both the 2021 and Post-2021 periods (Fig. 4). Furthermore, thinning treatments exhibited weaker suppression of seasonal NDVI gains during and after the heatdome, whereas control plots experienced prolonged reductions in NDVI responsiveness to seasonal progression (Fig. 5). These results suggest that thinning promoted stronger NDVI recovery after the heatdome compared to controls (Case et al. 2023, Loverin et al. 2024).

The differing trajectories of NDVI in the restoration treatments also highlighted potential differences in responses according to stand age. Specifically, younger stands (control and treatment) exhibited a smaller magnitude of NDVI suppression compared to the old-growth sites. The unmanaged old-growth forests likely still have higher canopy complexity than both the thinned and control stands, which rendered them more susceptible to the heatdome event (Chamberlain et al. 2021). Typical of intensively managed forests, Ellsworth consists of forests that were planted to be dense while still permitting radial tree growth for efficient timber production, leading to a relatively high canopy overlap in younger forests which is likely much higher than during unmanaged succession in these forests.

### Complex considerations in large scale forest restoration treatments

In all, our work adds to other studies in this system illustrating challenges that forest managers face in balancing multiple objectives in a changing environment. The initial study was designed to address whether thinning could help achieve structural complexity more rapidly, with our recent studies adding insight into how these treatments affect microclimate buffering and responses to extreme events (John et al. 2024). We found that dense, unthinned stands (old-growth and control) exhibited the strongest NDVI suppression during and after 2021, while thinned stands maintained more stable seasonal NDVI and showed quicker recovery i.e., higher canopy functional resilience despite potentially providing lower instantaneous thermal buffering. This “split result” suggests that residual canopy cover does buffer near-surface temperatures but that benefits can be limited under extreme conditions, so microclimate protection may not fully prevent physiological stress signals in the canopy (e.g., NDVI declines) (Brackett et al. 2024). More broadly, forest microclimates and structure jointly mediate organismal and canopy responses, with composition and canopy traits shaping when/where buffering fails during hot, dry spells (Zellweger et al. 2020).

During the June 2021 PNW heatdome, understories were ∼3 °C cooler than a nearby clear-cut and ∼4 °C cooler than the regional macroclimate, with stronger cooling under denser canopies—demonstrating that buffering persisted even under extreme heat, although unevenly across structures (John et al. 2024). This aligns with global syntheses showing that closed canopies systematically offset extreme heat in forest microclimates (and that offsets can intensify under extremes), and that structural attributes govern the magnitude of that effect (Zellweger et al. 2019, De Frenne et al. 2019). But, targeted thinning to restore old-growth structure does not substantially diminish microclimate buffering—supporting the idea that restoration goals and microclimate protection need not be in inherent conflict (Pradhan et al. 2023).

The canopy stress signal likely induced by the 2021 heatdome was persistent across forest types but did differ in its impacts according to treatment (Fig.4 and Fig.5). The clearest suppression occurred in control (dense) and thinned canopy structures, supporting the view that forest architecture mediates sensitivity to short-term climatic extremes. These findings highlight the utility of NDVI as a monitoring tool for detecting forest stress and assessing resilience to extreme heat events(Kulesza and Hościło 2023, Matiza et al. 2024).

Taken together, our results suggest that there is a management trade-off: thinning can advance structural development toward old-growth conditions and can enhance resilience and recovery following extreme heat events, but it slightly reduces immediate thermal buffering compared with the densest canopies during a heatwave. Managers can balance this tradeoff by designing spatially heterogeneous treatments—retaining dense patches as thermal refugia while thinning elsewhere to bolster canopy function and recovery potential—an approach that is consistent with the climate-refugia framework in restoration practice (Still et al. 2023, Pradhan et al. 2023). Where available, structure-aware mapping (e.g., LiDAR-derived canopy metrics) can target where buffering is strongest and where resilience gains from thinning are most likely, improving placement of treatments under a warming, more extreme climate regime (Au et al. 2022, Gril et al. 2023, John et al. 2024).

## Conclusions

As the frequency and duration of episodic extreme heat events continue to increase (IPCC 2022), the ability of forests to buffer heat stress is increasingly critical. Our study quantifies the impact of the 2021 Pacific Northwest (PNW) heatdome on forest canopy greenness and reveals that forests experienced significant stress during this extreme event. Differences in NDVI suppression between forested and harvested areas were substantial, suggesting that forest cover can moderate heat stress for terrestrial ecosystems. However, our results demonstrate that not all forests provide the same degree of protection. While thinned second-growth forests exhibited more stable NDVI trajectories and faster recovery following the heatdome, unthinned second-growth and dense old-growth plots experienced more pronounced and prolonged suppression. These patterns highlight that although denser canopies can initially buffer environmental extremes, their ability to withstand cumulative heat stress may be limited under sustained thermal events On a more practical level, we believe that a better understanding of the relationship between forest cover and buffering during extreme heat events could provide insight into best practices for forest restoration and management in a world characterized by warmer temperatures and more frequent extreme events. In our study system, canopy cover was manipulated through experimental thinning designed to accelerate the restoration of old-growth forest structure. Our results suggest that moderate thinning, which reduced canopy density, may have buffered forests against extreme heat stress. Thinned plots exhibited higher NDVI values and more stable seasonal trajectories during and after the 2021 heatdome compared to unthinned control plots, indicating that selective canopy reduction can promote greater forest resilience under acute climatic disturbances. Thus, it is possible that as these heatdome events increase in frequency and magnitude given warming trends (Guerreiro et al. 2018, Clarke et al. 2022), their increased negative impacts on thinned forests (e.g. potentially forest regeneration) could interfere with conservation and restoration goals. However, the impacts of climate change on these thinned forests might differ between the short-term extreme temperatures (explored in this study) vs. average seasonal temperatures. Short term temperature extremes are less buffered in thinned forest stands but management through thinning did not alter the thermal buffering capacity in the same watershed over longer time periods (Pradhan et al. 2023, John et al. 2024). Collectively, these three studies highlight the complexity of decision-making for conservation and the importance of examining a landscape from different perspectives. If this is possible, managers may be better able to balance restoration goals achieved through density reduction while maintaining resilience to climate change.

## Author contributions

J.H.R.L, K.P., M.C., A.K.E. and A.J. designed the study and collected the data. A.J. and K.P. analyzed the data with assistance from A.K.E. and J.H.R.L. A.J. and K.P. developed the first draft of the manuscript. All the authors contributed to reviewing and editing the draft.

## Acknowledgments

We acknowledge that some of this research was conducted in ancient lands of Lower Chinook people, the land which touches the shared waters of all tribes and bands within the Confederated Tribes of Grand Ronde, Chinook, and Confederated Tribes of Siletz Indians. We are grateful to the Planet Labs for providing the satellite imagery and computational support. We also thank The Nature Conservancy for the opportunity to conduct world-class climate research in their experimental forest.

## Data Availability Statement

The data that support the findings of this study are openly available in Zenodo at https://doi.org/10.5281/zenodo.18204628

John, A., K. Pradhan, M. J. Case, A. K. Ettinger, and J. Hille Ris Lambers. 2024. Data and code for assessing heatdome effects on forest dynamics in the Pacific Northwest using Planet imagery. Zenodo. https://doi.org/10.5281/zenodo.18204628

## Appendix A

**Table A1:** Summary of marginal effects of day of year (DOY) on NDVI from the GAM model, across forest treatment × period groups. The numbers (Min Slope, Max Slope) represent the range of slopes (rate of NDVI change per unit of DOY) estimated by the GAM smooths. Positive values mean NDVI is increasing with time (greening), and negative values means NDVI decreasing with time (suppression).

| Treatment | Period | Mean Slope | Min Slope | Max Slope | Lower 95% | Upper 95% |
| --- | --- | --- | --- | --- | --- | --- |
| OLD_GROWTH | Pre2021 | 0.00006 | -0.00009 | 0.00019 | 0.00014 | -0.00002 |
| OLD_GROWTH | 2021 | -0.00076 | -0.00077 | -0.00075 | -0.00055 | -0.00097 |
| OLD_GROWTH | Post2021 | 0.00148 | -0.00339 | 0.00665 | 0.00163 | 0.00132 |
| CON | Pre2021 | 0.00002 | -0.00059 | 0.00049 | 0.00008 | -0.00004 |
| CON | 2021 | -0.00088 | -0.00111 | -0.00062 | -0.00063 | -0.00114 |
| CON | Post2021 | 0.00055 | -0.00436 | 0.00598 | 0.00059 | 0.00051 |
| THIN | Pre2021 | 0.00015 | -0.00021 | 0.00044 | 0.00020 | 0.00010 |
| THIN | 2021 | -0.00035 | -0.00079 | -0.00002 | -0.00009 | -0.00062 |
| THIN | Post2021 | 0.00054 | -0.00472 | 0.00632 | 0.00058 | 0.00049 |
| CLEARCUT | Pre2021 | 0.00033 | -0.00032 | 0.00103 | 0.00050 | 0.00017 |
| CLEARCUT | 2021 | 0.00032 | -0.00353 | 0.00375 | 0.00144 | -0.00080 |
| CLEARCUT | Post2021 | 0.00034 | -0.01478 | 0.02225 | 0.00055 | 0.00012 |

**Figure A1.**
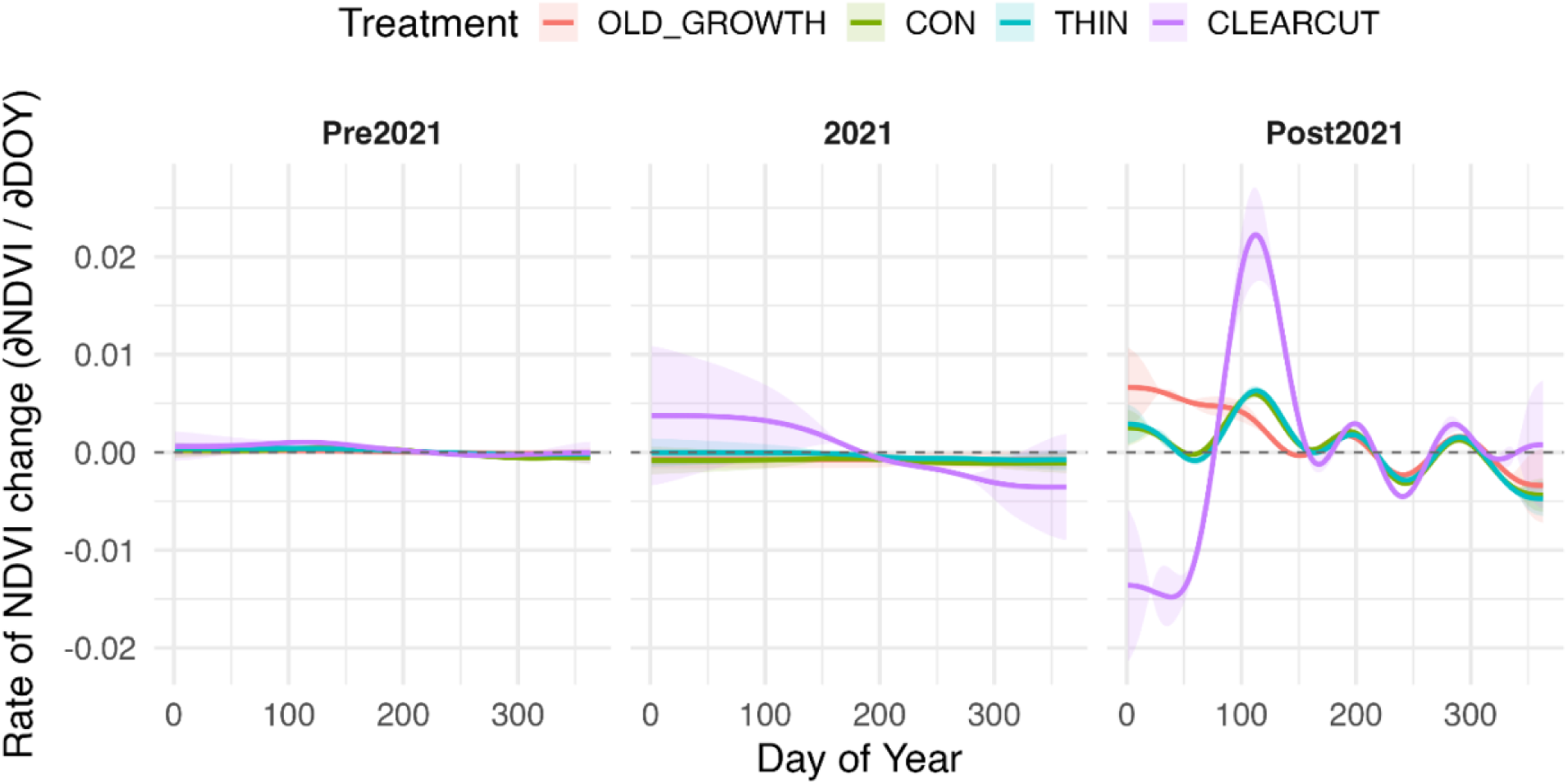
Marginal effects of day of year (DOY) on NDVI across forest treatments and periods from the GAM analysis. Marginal effect curves represent the rate of NDVI change relative to the day of year across three temporal periods (Pre-2021, 2021, Post-2021) within each treatment (OLD_GROWTH, CON, THIN, CLEARCUT). Lines represent fitted marginal effects and shaded ribbons indicate 95% confidence intervals. During 2021, particularly in CON plots, marginal effects were negative (suppressed), reflecting reduced seasonal green-up during the heatdome. In contrast, THIN plots maintained relatively stable and positive marginal effects across all periods, suggesting greater resilience to heat stress.

**Table A2:**
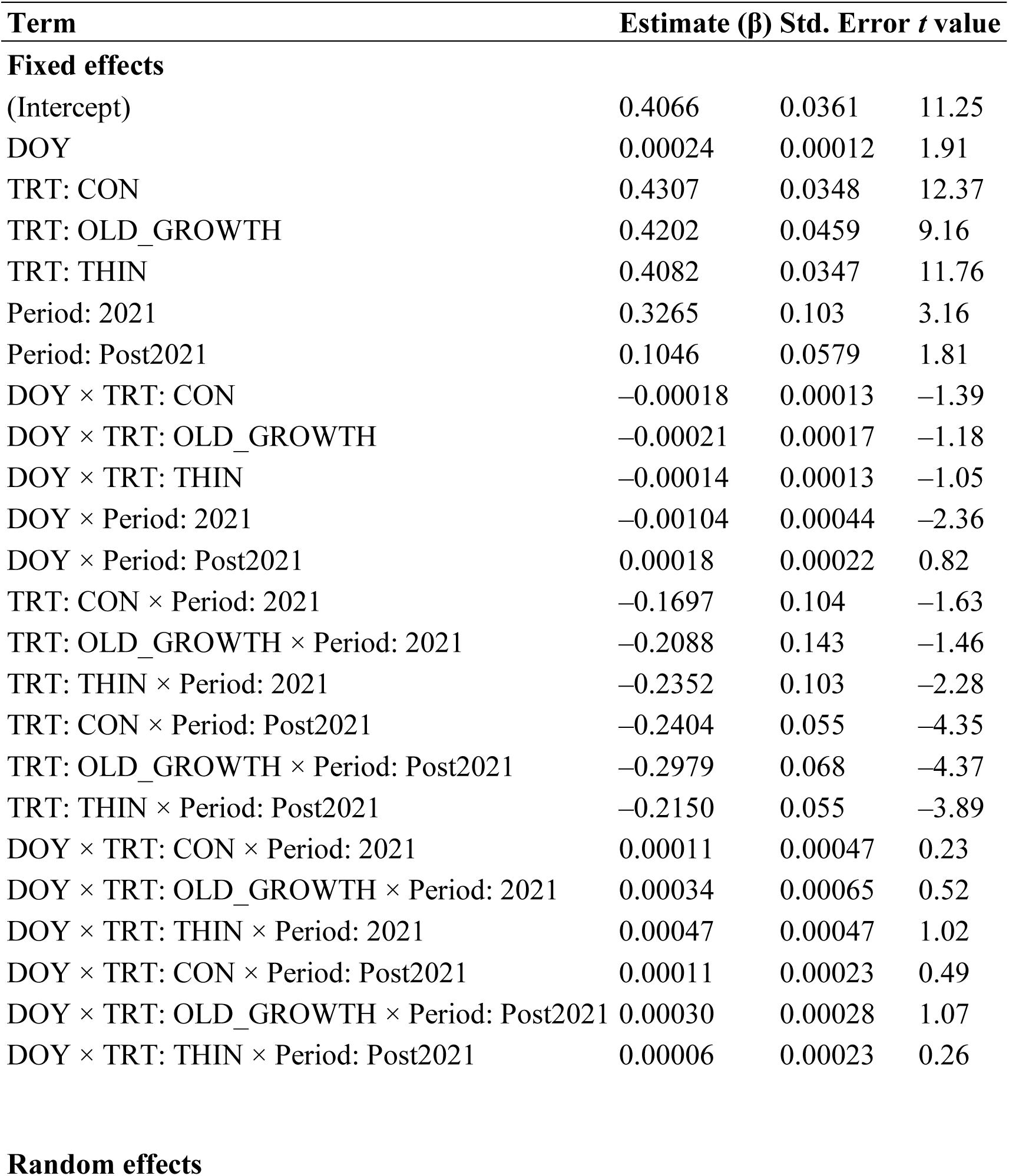
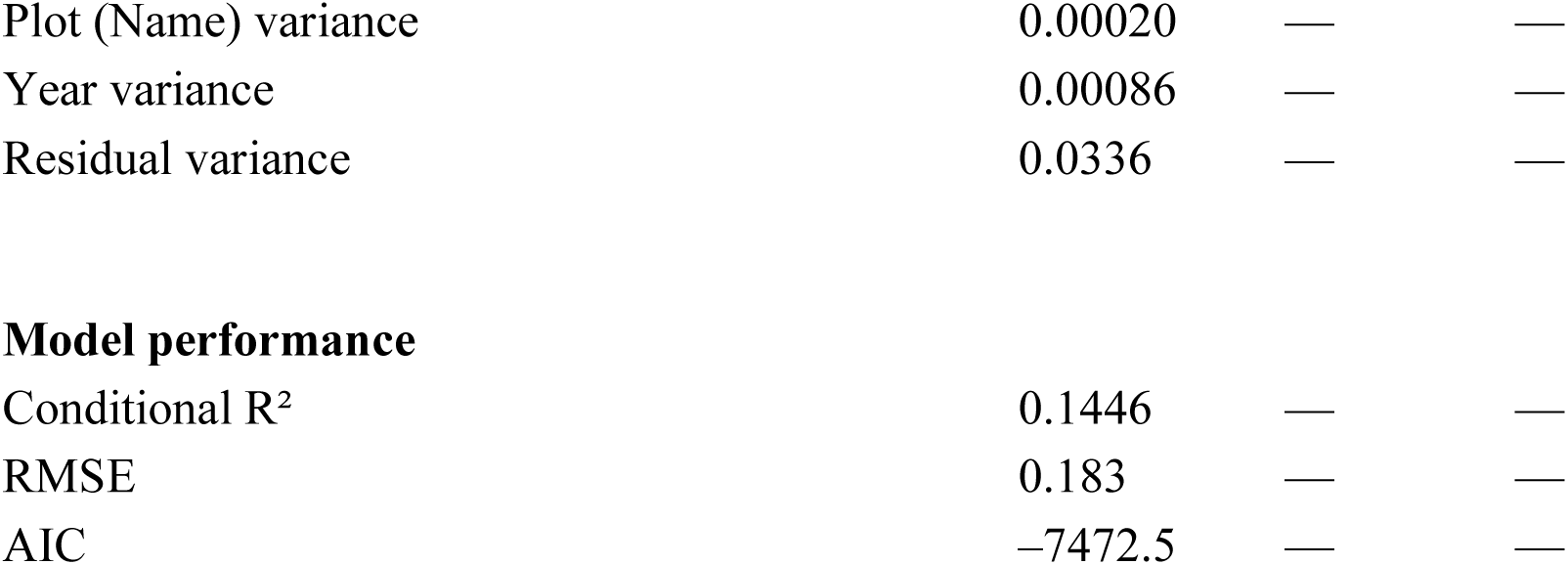
Full LMM results for NDVI ∼ DOY × Treatment × Period.

| Term | Estimate (β) | Std. Error | t value |
| --- | --- | --- | --- |
| **Fixed effects** |  |  |  |
| (Intercept) | 0.4066 | 0.0361 | 11.25 |
| DOY | 0.00024 | 0.00012 | 1.91 |
| TRT: CON | 0.4307 | 0.0348 | 12.37 |
| TRT: OLD_GROWTH | 0.4202 | 0.0459 | 9.16 |
| TRT: THIN | 0.4082 | 0.0347 | 11.76 |
| Period: 2021 | 0.3265 | 0.103 | 3.16 |
| Period: Post2021 | 0.1046 | 0.0579 | 1.81 |
| DOY × TRT: CON | −0.00018 | 0.00013 | −1.39 |
| DOY × TRT: OLD_GROWTH | −0.00021 | 0.00017 | −1.18 |
| DOY × TRT: THIN | −0.00014 | 0.00013 | −1.05 |
| DOY × Period: 2021 | −0.00104 | 0.00044 | −2.36 |
| DOY × Period: Post2021 | 0.00018 | 0.00022 | 0.82 |
| TRT: CON × Period: 2021 | −0.1697 | 0.104 | −1.63 |
| TRT: OLD_GROWTH × Period: 2021 | −0.2088 | 0.143 | −1.46 |
| TRT: THIN × Period: 2021 | −0.2352 | 0.103 | −2.28 |
| TRT: CON × Period: Post2021 | −0.2404 | 0.055 | −4.35 |
| TRT: OLD_GROWTH × Period: Post2021 | −0.2979 | 0.068 | −4.37 |
| TRT: THIN × Period: Post2021 | −0.2150 | 0.055 | −3.89 |
| DOY × TRT: CON × Period: 2021 | 0.00011 | 0.00047 | 0.23 |
| DOY × TRT: OLD_GROWTH × Period: 2021 | 0.00034 | 0.00065 | 0.52 |
| DOY × TRT: THIN × Period: 2021 | 0.00047 | 0.00047 | 1.02 |
| DOY × TRT: CON × Period: Post2021 | 0.00011 | 0.00023 | 0.49 |
| DOY × TRT: OLD_GROWTH × Period: Post2021 | 0.00030 | 0.00028 | 1.07 |
| DOY × TRT: THIN × Period: Post2021 | 0.00006 | 0.00023 | 0.26 |

Random effects
| Term | Estimate ($\beta$) | Std. Error | *t* value |
| --- | --- | --- | --- |
| Plot (Name) variance | 0.00020 | — | — |
| Year variance | 0.00086 | — | — |
| Residual variance | 0.0336 | — | — |
| **Model performance** |  |  |  |
| Conditional R<sup>2</sup> | 0.1446 | — | — |
| RMSE | 0.183 | — | — |
| AIC | −7472.5 | — | — |

**Figure A2.**
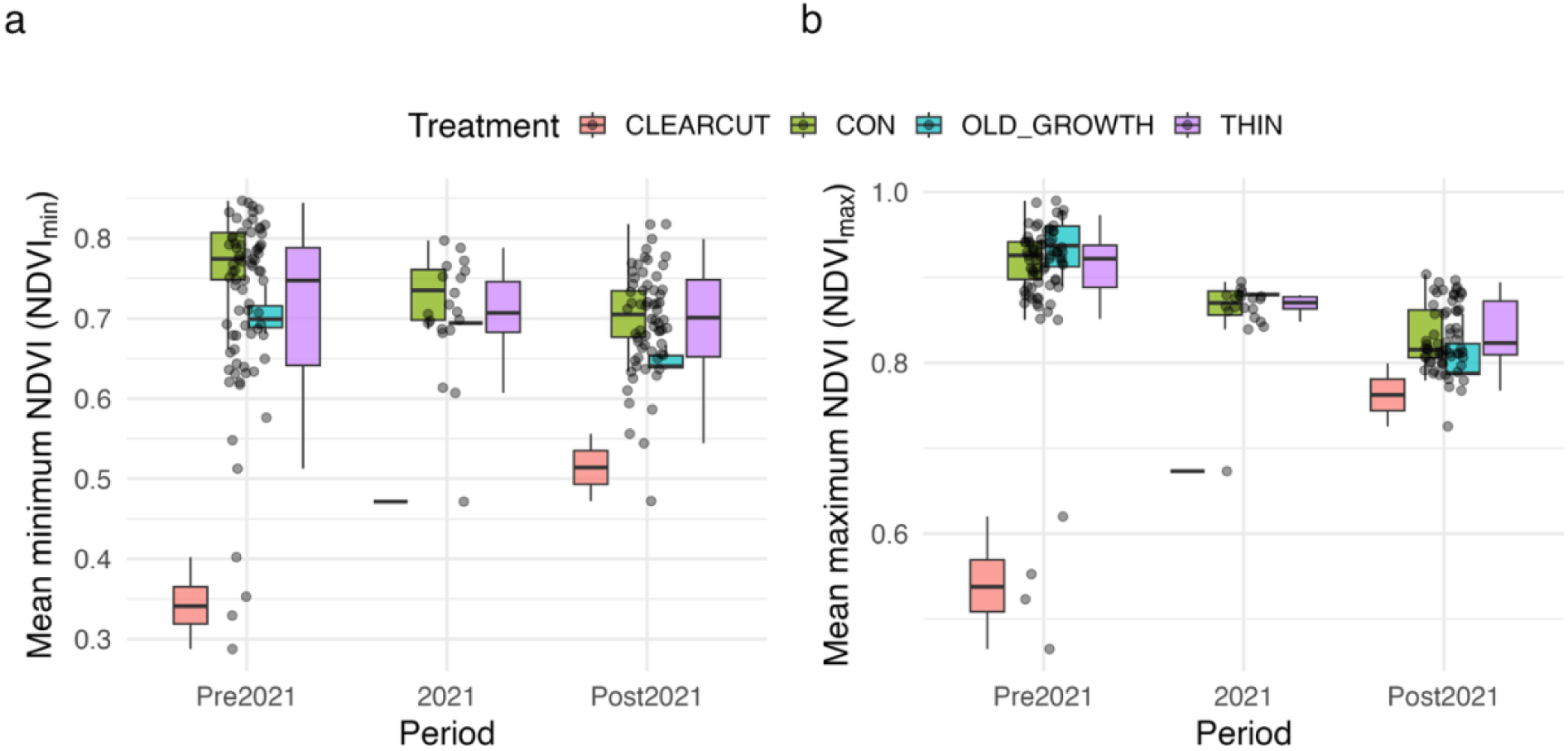
Minimum and maximum NDVI (NDVIₘᵢₙ, NDVIₘₐₓ) across forest treatments and temporal periods. (a) NDVIₘᵢₙ reflects baseline canopy greenness and structural persistence through the season, whereas (b) NDVIₘₐₓ indicates peak canopy greenness. Both metrics declined during the 2021 heatdome year, particularly in OLD_GROWTH and control plots, consistent with heat stress–induced canopy suppression. Post-2021, NDVIₘᵢₙ showed gradual recovery in thinned and clearcut plots, suggesting improving structural greenness, while NDVIₘₐₓ remained below pre-2021 levels in old-growth and clearcut stands, indicating incomplete recovery of canopy photosynthetic potential.

## Appendix B

**Figure B1.**
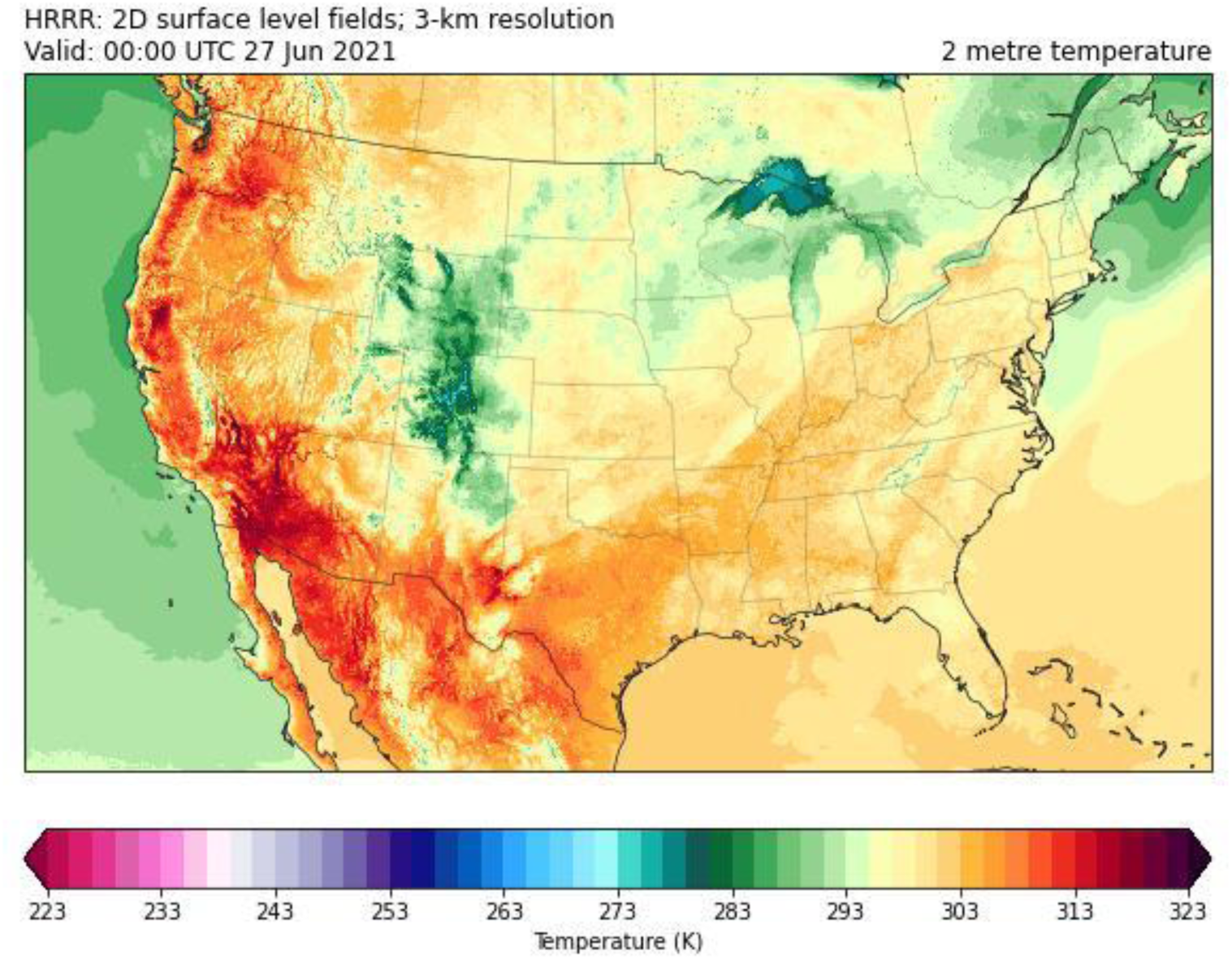
Spatial extent of the late-June 2021 North American heatdome captured by the High-Resolution Rapid Refresh (HRRR) model. Shown are 2-m air temperatures (K) from 3-km resolution surface fields on 27 June 2021, showing extreme and spatially warming across the western United States, including the Pacific Northwest

